# Interactive, visual simulation of a spatio-temporal model of gas exchange in the human alveolus

**DOI:** 10.1101/2021.09.15.460416

**Authors:** Kerstin Schmid, Andreas Knote, Alexander Mück, Keram Pfeiffer, Sebastian von Mammen, Sabine C. Fischer

## Abstract

In interdisciplinary fields such as systems biology, close collaboration between experimentalists and theorists is crucial for the success of a project. Theoretical modeling in physiology usually describes complex systems with many interdependencies. On one hand, these models have to be grounded on experimental data. On the other hand, experimenters must be able to penetrate the model in its dependencies in order to correctly interpret the results in the physiological context. When theorists and experimenters collaborate, communicating results and ideas is sometimes challenging. We promote interactive, visual simulations as an engaging way to communicate theoretical models in physiology and to thereby advance our understanding of the process of interest. We defined a new spatio-temporal model for gas exchange in the human alveolus and implemented it in an interactive simulation software named *Alvin*. In *Alvin*, the course of the simulation can be traced in a three-dimensional rendering of an alveolus and dynamic plots. The user can interact by configuring essential model parameters. *Alvin* allows to run and compare multiple simulation instances simultaneously. The mathematical model was developed with the aim of visualization and the simulation software was engineered based on a requirements analysis. Our work resulted in an integrative gas exchange model and an interactive application that exceed the current standards. We exemplified the use of *Alvin* for research by identifying unknown dependencies in published experimental data. Employing a detailed questionnaire, we showed the benefits of *Alvin* for education. We postulate that interactive, visual simulation of theoretical models, as we have implemented with *Alvin* on respiratory processes in the alveolus, can be of great help for communication between specialists and thereby advancing research.

## 1 Introduction

Systems biology is a highly interdisciplinary research field that integrates theoretical modeling and experimental data (Gavaghan et al., 2006). A key component of projects with valuable scientific progress is close cooperation between experimentalists and theorists (Byrne et al., 2006; Drubin and Oster, 2010; Welsh et al., 2006). However, this entails certain challenges. Different ways of thinking and terminologies or jargon often hinder communication between the disciplines. Ongoing efforts to bridge the gap include educational reviews (e.g. (Sharpe, 2017; Fischer, 2019)), summer schools, special research programs (https://www.newton.ac.uk/event/cgp/) and large multi-laboratory initiatives such as the Virtual Physiological Human (Viceconti et al., 2008) or The Virtual Brain (https://www.thevirtualbrain.org). Key components of these approaches are informative visualizations and the possibility of hands-on experience.

We consider communicating results of mathematical modeling in physiology. In publications, models are usually presented as follows (Mogilner et al., 2011): The model definition is given in terms of mathematical equations, occasionally supported by schematic diagrams describing the model structure. For the corresponding simulations, all parameter values are listed and the output is visualized in graphs and compared with experimental data, where appropriate. When modeling spatial structures and processes, the simulation output is presented in still images or, if possible, animations (Chao, 2003; Saber and Heydari, 2012; Lin et al., 2004). As an alternative for the communication of state of the art theoretical models, we promote interactive, visual simulation. Previous approaches include computer-aided diagnosis software (Xiong et al., 2017; Conover et al., 2018) or systems for medical education (Jacob et al., 2012; Jamniczky et al., 2012; Costabile, 2021). We focus on the human lung. Existing interactive systems for teaching in this field address respiratory mechanics (Kuebler et al., 2007; Warliah et al., 2012) or gas exchange (Kapitan, 2008). All above systems for teaching convey established educational content. They have not been intended to advance the current state of research.

In contrast, (Winkler et al., 1995) argue that their interactive system has great utility beyond its educational use. They have developed an application that provides an interactive interface with a simulation of a multi-compartment model. Ventilation mechanics, gas transport, gas mixing and gas exchange are considered. However, the actual process of gas exchange, the key functionality of the human lung, remains as abstract as the site where it occurs. We thus focused on the smallest functional unit of the lung - the alveolus - and developed an interactive, visual simulation of gas exchange. We refined and combined existing models (Weibel et al., 1993; Dash et al., 2016) to cover the complete transport of oxygen into hemoglobin. The resulting model provided the computational core for an interactive simulation software named *Alvin. Alvin* facilitates investigations of relationships between morphological and physiological factors and the course of gas exchange. The software enables systematic investigations of our model with respect to experimental data. The intuitive usability of *Alvin* allows its integration in research and teaching. As an exemplary use case in research, we present a plausibility check of pulmonary diffusion capacity measurements. Concerning the applicability of *Alvin* in teaching, we present the details of its integration into a digital physiology lab course for undergraduate students and the results of a corresponding survey among its participants. The software is available for download at https://go.uniwue.de/alvin.

Particular about our work is the development of the mathematical model with the aim of visualization in combination with the requirements-based engineering of the simulation software. This resulted in an advanced gas exchange model and an interactive application that exceed the existing standard. Specifically, design features as the ability to run and compare multiple simulation instances at the same time and the combination of providing parameter value presets as well as allowing parameter configurations by the user are key contributions to the field. This results in an educationally valuable application that also allows revealing unknown underlying assumptions of results presented in the literature. Taken together, our work demonstrates that an interactive, visual simulation can improve the understanding of gas exchange in the human lung, for both researchers and students.

## 2 Integrative Alveolar Gas Exchange Model

The human lung consists of progressively branching bronchi and bronchioles, and blood vessels follow this structure (Hsia et al., 2016). The respiratory zone begins where the first alveoli adjoin the bronchioles (Haefeli-Bleuer and Weibel, 1988). Alveoli are hollow protrusions that have a large surface area and a thin tissue barrier. They are surrounded by a dense network of fine capillaries (Weibel and Gomez, 1962). Within an alveolus, inhaled air passes through the cavity and gas exchange with the capillary blood takes place through the tissue barrier (Weibel, 2009). An alveolus thus represents the smallest functional unit of the lung.

### 2.1 Model

The process of gas exchange in an alveolus can be divided into two sequential steps (Roughton and Forster, 1957): 1. The diffusion of oxygen through the tissue barrier into the blood and red blood cells and 2. its binding to hemoglobin (Hb). Gas exchange leads to oxygen (O_2_) and carbon dioxide (CO_2_) pressure gradients inside the blood along capillaries surrounding the alveolus. In a healthy individual, blood enters this area with a low partial pressure of oxygen (pO_2_) and a high partial pressure of carbon dioxide (pCO_2_). Diffusion of O_2_ from the alveolus into the capillary and of CO_2_ out of the capillary into the alveolus gradually increases pO_2_ and decreases pCO_2_ until the distribution of gases reaches equilibrium (Powers and Dhamoon, 2019). Hence, the course of pressure gradients depends on the efficiency of gas diffusion and the blood flow velocity. To map O_2_ and CO_2_ pressure gradients in our model, a representative capillary was divided into subsections of equal size (Figure 1). Oxygen diffusion from the alveolar space into the different sections is calculated successively starting with the first section. Here, blood enters with a preset pO_2_. The oxygen flow *ν* across the barrier as a function of morphological parameters and the pressure gradient ΔpO_2_ between air and blood is calculated based on Fick’s diffusion law (Weibel et al., 1993), resulting in

**Figure 1:**
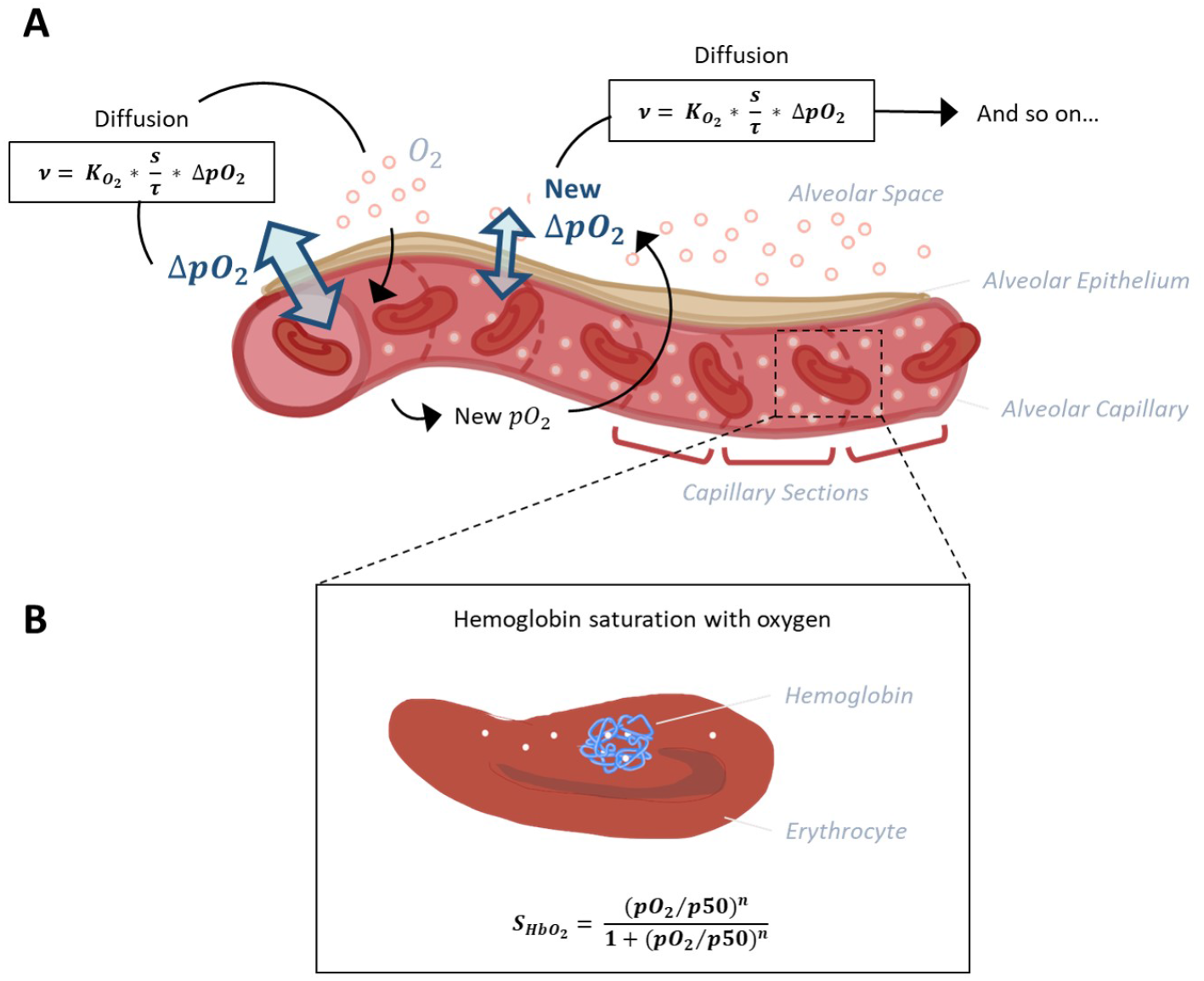
Schematic representation of the model capillary with erythrocytes, separated from alveolar space by a single cell layer of alveolar epithelium. **(A)** In order to reconstruct O_2_ and CO_2_ pressure gradients along the capillary, it is divided into sections of equal size. The pressure gradient between alveolar space and blood (ΔpO_2_) and the resulting flow of oxygen along this gradient is calculated for each section subsequently, as oxygen flow into one section affects pO_2_ and thus ΔpO_2_ of the next section. Calculation of oxygen diffusion depending on ΔpO_2_ is based on Fick’s law (Weibel et al., 1993). **(B)** According to the pO_2_ and pCO_2_ gradients along the capillary sections determined in step 1, hemoglobin oxygen saturation 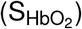 is calculated for each section. The corresponding Hill equation has been defined and fitted to experimental data (Dash et al., 2016).

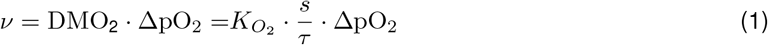

The membrane diffusing capacity for oxygen DMO_2_ comprises the ratio between surface area *s* and barrier thickness *τ* multiplied by the permeability coefficient 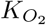. The amount of oxygen reaching the next capillary section depends on the blood flow velocity. The incoming oxygen affects the pO_2_ of blood in that section and so on.

The quantity of CO_2_ diffusing out of the capillary and into the alveolus is determined via the respiratory exchange ratio from the quantity of oxygen that is taken up by the blood. The respiratory exchange ratio is defined as the amount of CO_2_ produced divided by the amount of O_2_ consumed. This ratio is assessed by analyzing exhaled air in comparison with the environmental air and its average value for the human diet is around 0.82 (Sharma et al., 2020).

The part of our mathematical model describing the binding of O_2_ and CO_2_ to hemoglobin was adopted from (Dash et al., 2016), such that

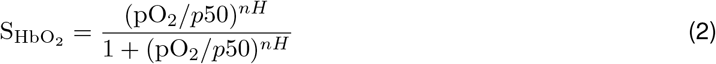

Hemoglobin oxygen saturation 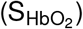 is expressed as a Hill function depending on pO_2_, the Hill coefficient *nH* and *p*50, the value of pO_2_ at which hemoglobin is 50 % saturated with O_2_. The parameter *nH*, in turn, depends on pO_2_. Polynomial expressions describe the dependence of *p*50 on pCO_2_ in the blood, blood temperature, the pH inside erythrocytes (pH_rbc_) and concentration of the organic phosphate 2,3-bisphosphoglycerate ([2,3]-DPG). These dependencies have been described and fitted to several experimental data sets (Dash et al., 2016). In our model, 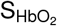 is determined for each section according to the pO_2_ and pCO_2_ gradients along the capillary sections obtained from step 1.

Together, we obtain a model for the complete process of oxygen transport from inhaled air into hemoglobin in the blood with spatio-temporal resolution. All parameters essential for the model and their default values were collected from the literature and represent a normal, healthy condition (Table 1).

**Table 1:** Model parameters and their default values. Values of morphological and physiological parameters of the gas exchange model were collected from literature. All values given are mean values referring to a single alveolus.

| Parameter | Unit | Default Value | Reference | Value Range |
| --- | --- | --- | --- | --- |
| Alveolar pO<sub>2</sub> | mmHg | 100 | (Sharma et al., 2020) | 1 - 150 |
| Blood pO<sub>2</sub> | mmHg | 40 | (Dash et al., 2016) | 1 - 150 |
| Alveolar pCO<sub>2</sub> | mmHg | 40 | (Sharma et al., 2020) | 1 - 150 |
| Blood pCO<sub>2</sub> | mmHg | 45 | (Dash et al., 2016) | 1 - 150 |
| Surface area | μm<sup>2</sup> | 121000 | (Mercer et al., 1994) | 0 - 210000 |
| Thickness of tissue barrier | μm | 1.11 | (Gehr et al., 1978; Weibel et al., 1993) | 0.1 - 0.3 |
| Blood flow velocity | mm/s | 1 | Abstracted from: (Weibel et al., 1993; Petersson and Glenny, 2014) | 0.01 - 2 |
| Blood volume | μm<sup>3</sup> | 404000 (50 % "capillary recruitment") | Abstracted from: (Gehr et al., 1978; Ochs et al., 2004; Okada et al., 1992) | 1 - 880000 |
| Blood temperature | °C | 37 | (Dash et al., 2016) | 20 - 44 |
| Erythrocyte pH (pH<sub>rbc</sub>) |  | 7.24 | (Dash et al., 2016) | 5.8 - 8.2 |
| Concentration of [2,3]-DPG | mM | 4.65 | (Dash et al., 2016) | 1 - 10 |
| Capillary length | μm | 500 | (Weibel et al., 1993) | *not adjustable |
| Capillary volume | μm<sup>3</sup> | 808000 | (Ochs et al., 2004; Gehr et al., 1978) | *not adjustable |
| Capillary radius | μm | 3.15 | (Mühlfeld et al., 2010) | *not adjustable |
| Number of capillaries |  | 52 | Calculated from capillary volume, radius and length | *not adjustable |

### 2.2 Model validation

In a first step of model validation, we analysed whether the two sub models from step 1 and step 2 had been sensibly adapted from the literature. In our model, oxygen diffusion is estimated for a single alveolus with a surface area of 121000 µm^2^. Other parameters affecting DMO_2_ (namely tissue barrier thickness and permeability coefficient, see Equation 1) were adopted without change. DMO_2_ of the whole lung in relation to body weight (bw) was estimated as 0.079 mL/(s · mmHg · kg) (Weibel et al., 1993). To compare our model result (DMO_2_^(model)^ = 6 · 10^*−*9^ mL/(s · mmHg)) with Weibel’s estimate, it needs to be extrapolated to the organ scale. Multiplying DMO_2_^(model)^ by the number of alveoli in the human lung (480 ·10^6^ (Ochs et al., 2004)) results in a DMO_2_^(model, extrapolated)^ of 2.88 mL/(s · mmHg). This value is distinctly lower than the DMO_2_ estimated by Weibel et al., assuming a standard body weight of 70 kg: DMO_2_^(Weibel, bw 70 kg)^ = 5.53 mL/(s · mmHg). This estimate has been based on morphometric studies in fully inflated, fluid-filled lungs (Weibel et al., 1993). It is recognized that in an air-filled lung, however, only about 60-70% of the alveolar surface is exposed to air (Gil et al., 1979; Bachofen et al., 1987). The default value for surface area in our model was taken from studies on perfusion-fixed, air-filled lungs (Mercer et al., 1994). Hence, our combination of parameter values for the surface area of a single alveolus (Mercer et al., 1994) and the number of alveoli in the human lung (Ochs et al., 2004) produce a result that falls short of the previous estimate. However, the discrepancy is explained by known differences in the morphometric methods used. We deliberately chose the surface value from the study on an air-filled lung to be as close as possible to the *in vivo* situation. The sub model describing hemoglobin oxygen saturation was adopted from the literature (Dash et al., 2016) without further modifications. Hb-O_2_ dissociation curves across the different parameter ranges from this publication (Figure 4 E-H in (Dash et al., 2016)) were recreated and indicate a correct implementation of the model (Figure 2).

**Figure 2:**
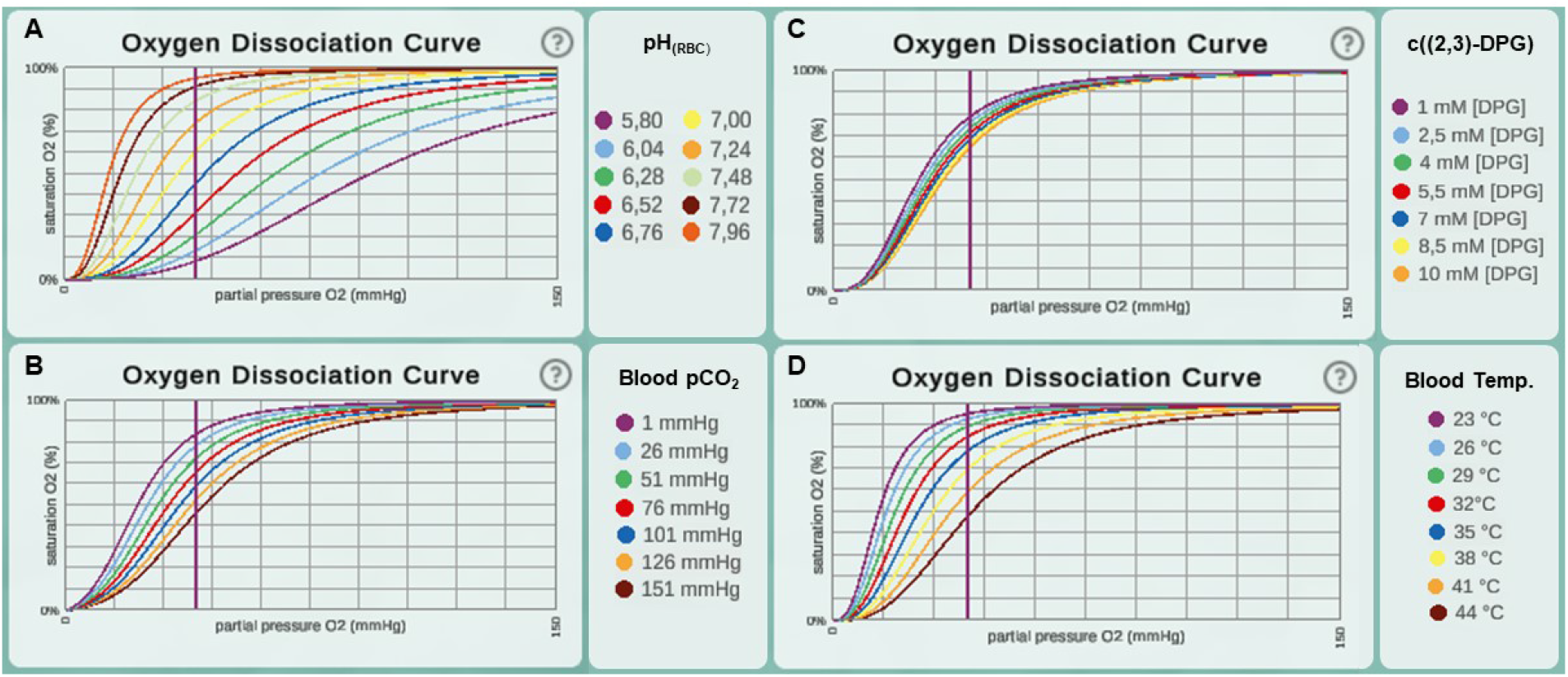
Oxygen dissociation curves for different ranges of parameter values from the original paper (Dash et al., 2016). This includes value ranges for the parameters **(A)** pH in erythrocytes (pH_rbc_), **(B)** blood pCO_2_, **(C)** concentration of [2,3]-DPG and **(D)** blood temperature.

In a second step, the complete integrative model was validated. We used published experimental data to validate our model. A key contribution of our model is the temporal and spatial resolution. Rather than determining mean values, oxygen partial pressure and saturation gradients along the alveolar capillary are generated. This allows validation of the model in a physiological context. For default parameter settings, 50 % of the oxygenation that blood undergoes during its transit along the alveolus is completed after 0.04 s (Figure 3). This measurement was performed for an increase in saturation from 81 % to 97 %, reaching the reaction half-time at 89 %. The corresponding measurement in mice is 0.037 s (Tabuchi et al., 2013) and it has been argued that there are only slight differences between species (Lindstedt, 1984). In summary, we showed that we have correctly adopted and sensibly modified the individual models. Our new integrative model provides results that are consistent with experimental data.

**Figure 3:**
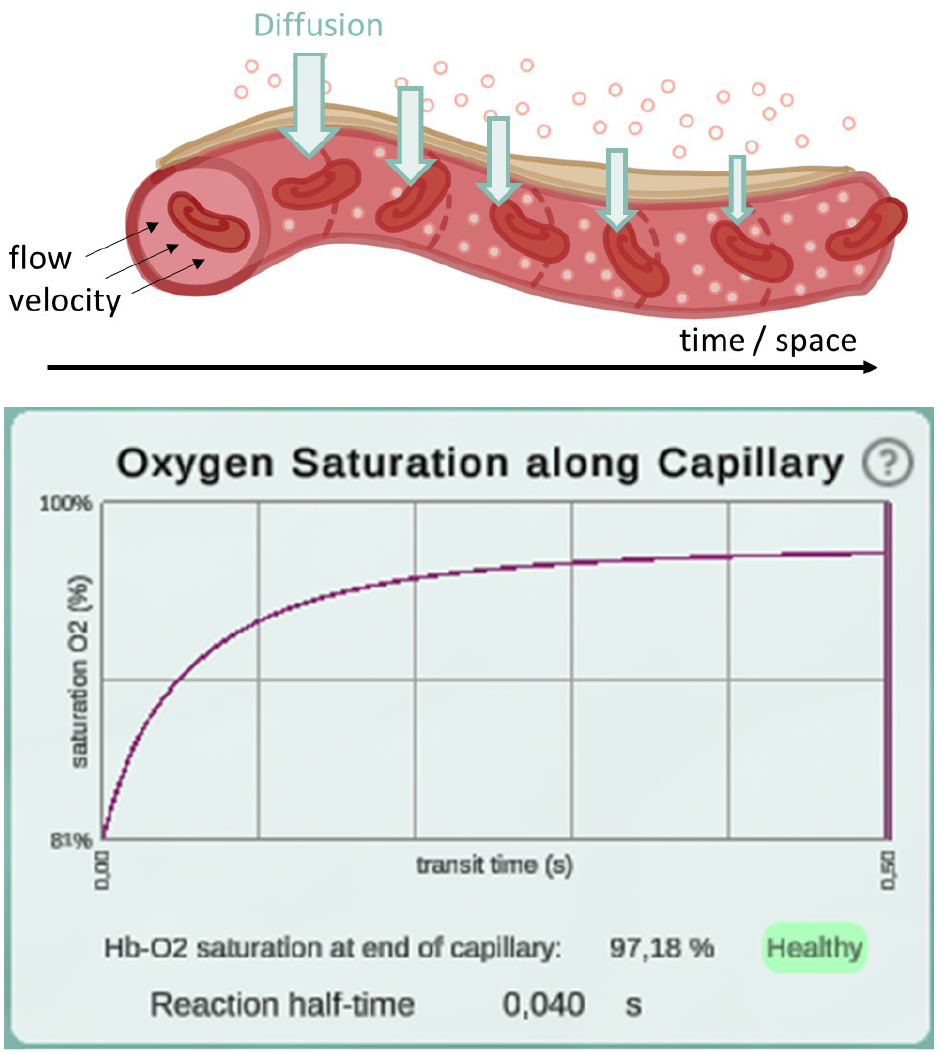
Illustration of the diffusion gradient along the model capillary (top) and a screenshot of the plot displaying oxygen saturation along capillary between 81 % and 97 % (bottom). This screenshot was taken from a simulation with pO_2_ values of 97 mmHg in the alveolar space and 46 mmHg in the deoxygenated blood. All other parameters remained at their default settings. Reaction half-time is defined as the time point at which 50 % of the oxygenation that blood undergoes during its transit along the alveolus is reached.

### 2.3 Model discussion

Our mathematical model was assembled from two existing sub models (Weibel et al., 1993; Dash et al., 2016). One sub model describes the diffusion rate of oxygen from the air into the blood depending on morphological properties (Weibel et al., 1993). In this preceding work, the lung has been defined simplistically as a single container of air and the partial pressure of oxygen in the blood has been considered constant. Some simplifications still exist in our new model. For example, the introduction of a breathing pattern was neglected: Partial pressure changes in alveolar space only occur when respective parameter values are modified by the user (suggests that O_2_ diffusing out of the alveolus is instantly replaced and CO_2_ diffusing into the alveolus is evacuated immediately). Also, blood flow was approximated as a continuous flow of a homogeneous plasma/ erythrocyte mixture. However, our new integrative model also features improvements compared to the original models. Instead of steady states, it provides information about oxygen transport over the continuous course of time. It has already been noted that a time-dependent modeling approach is better suited to reconstruct gas exchange in lung tissue than steady-state approaches (Sapoval et al., 2020). Accordingly, the temporal resolution is a valuable improvement to the model.

For validation, we compared reaction half-time results from our model with what has been reported in the literature (Tabuchi et al., 2013). Reaction half-time is defined as the time that elapses until 50 % of the oxygenation that blood undergoes during its transit along the alveolus is complete. We measured 40 ms with default parameter settings. Experimentally, a half-time of 37 ms has been determined in mice (Tabuchi et al., 2013). Corresponding theoretical predictions have been slightly lower at 18-32 ms. Tabuchi et al. argue that this discrepancy is due to the fact that the oxygenation process already takes place in the precapillary arterioles, but for the prediction only capillaries were considered. Since only capillaries are considered in *Alvin* as well, we may suspect that our value underestimates the *in vivo* human reaction half-time slightly.

In our model, capillaries are divided into an arbitrary number of sections. The finer grained this discretisation, i.e. the smaller the individual sections and the larger their number, the larger is the resolution of calculated gas dynamics and, thus, the resulting accuracy. However, as described in the following section, our model forms the basis of a visual simulation. With higher resolution, the computational demand grows, especially due to the three-dimensional rendering of the respective capillary sections. Therefore, we manually optimised this detail to maximise the accuracy without jeopardising the simulation’s interactivity.

## 3 Visualization and Interactivity: The *Alvin* Application

Interaction with content positively influences its conception (He et al., 2021; Pike et al., 2009) and helps to explore concepts. Having established the mathematical model, we developed the *Alvin* simulation software to support the conception and exploration of the gas exchange process in a single alveolus.

### 3.1 Requirements Analysis

We instigated the design process of the software by means of a requirements analysis. The target groups and the most important functionalities desired for this application were defined. Our core target group are scientists. *Alvin* communicates the state of the art in modeling gas exchange to colleagues in the context of research, or to undergraduate students in the context of teaching. The purpose of *Alvin* is not only to show final simulation results, but also to enable interactions with the model parameters during the simulation. This allows the exploration of the model and testing new hypotheses. This objective implies the following requirements:

Functional requirements (FR):

- FR1 The application should be able to run on common devices.
- FR2 Quantitative and qualitative simulation output should be provided.
- FR3 The simulation should respond directly to changes in parameter values by the user.
- FR4 The connection between parameter value changes and physiology should be provided.
- FR5 The connection between parameter value changes and output should be emphasized.
- FR6 Both healthy and disease conditions should be considered, and comparisons should be possible.

Non-functional requirements (NFR):

- NFR1 The application should allow the user to enter a flow state when exploring the model behaviour.
- NFR2 The overall look and feel of the application should be professional and clean as well as appealing and responsive.

### 3.2 Visualization

*Alvin* is a desktop-based application implemented in Unity. It is available for Windows, macOS and Linux (fulfills FR1). The user interface of *Alvin* consists of the following core components: a three-dimensional model of an alveolus illustrating the simulation process, a configuration menu for model parameter values and a panel displaying dynamic graphs (Figure 4). A key feature is the ability to run and compare multiple simulation instances at the same time.

**Figure 4:**
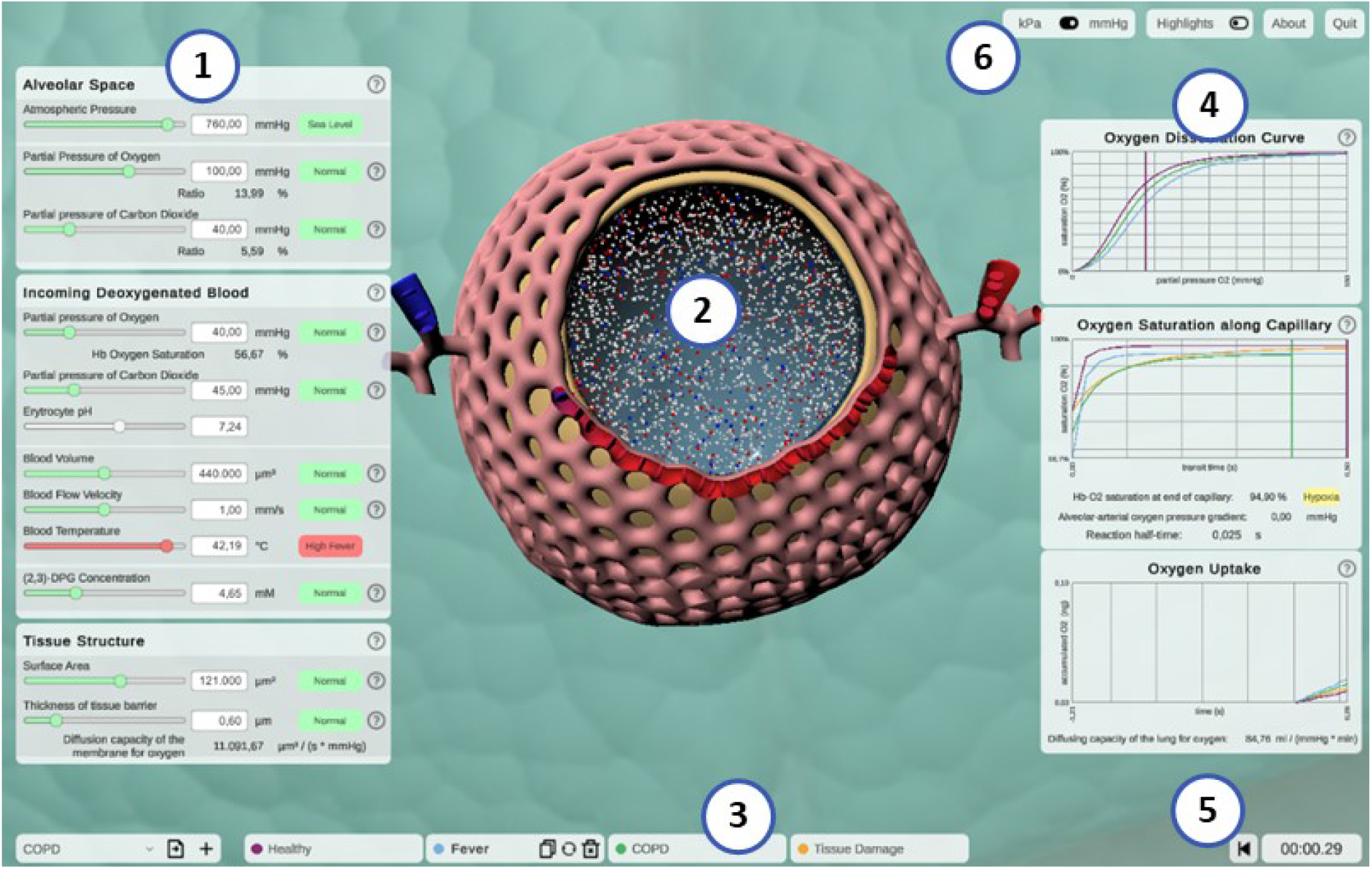
Screenshot of the interactive application *Alvin*. **(1)** Model parameters can be configured, grouped into categories. Colors and information text provide possible real-world interpretation of the values. **(2)** Animated simulation of an alveolus for the active parameter set provides visualization of the effect of the model parameter values. **(3)** To increase exploratory value, multiple simulation instances can be compared. **(4)** Quantitative simulation output is displayed with plots color-coded for each active instance of the simulation. **(5)** Simulation time is displayed and can be reset. **(6)** Utility functions and settings are available.

The animated, three-dimensional model of an alveolus illustrates the current state of the simulation (Figure 4, center, see also Section S.1.2 for further details). The alveolus is visually filled with small representations for air molecules, animated to signify Brownian motion. Each one is representing roughly 2 ·10^9^ molecules of oxygen (red spheres), carbon dioxide (blue spheres) or nitrogen (white spheres), respectively. Thickening or thinning of the tissue layer indicates value changes of the model parameter “thickness of tissue barrier”. Erythrocytes are animated and move along the cut-open capillary. The number of erythrocytes proportionally corresponds to a standard value of 5 · 10^6^ cells per µL blood (Pagana et al., 2019). Their relative position on this path is constantly tracked. Oxygen partial pressure (Equation 1) and hemoglobin oxygen saturation (Equation 2) gradients are calculated along the same path. This information is combined to color erythrocytes according to their oxygen saturation and to cumulatively total the amount of oxygen taken up by the erythrocytes over the course of the simulation (see Figure 4, graph “oxygen uptake”).

Hence, simulated gas exchange can be retraced by observing the amount of gas spheres crossing the tissue barrier from one side to the other and changes in capillary and erythrocyte coloring (FR2). Quantitative outcome of the simulation can be monitored on three different graphs (FR2)(Figure 4, right). They show hemoglobin oxygen saturation as a function of pO_2_ in the blood (oxygen dissociation curve), or of time (oxygen saturation along capillary). Finally, the total amount of oxygen taken up is tracked as a function of the time since the simulation was started or reset. Graphs of different simulation instances are indicated by their respective instance color.

### 3.3 Interactivity

The parameter panel (Figure 4, left) allows users to configure model parameter values. Changes in parameter values yield run-time updates in the 3D visualization and the quantitative graphs (FR3). A traffic light color code and keywords provide classification of the chosen parameter values with regard to their healthy or pathological ranges (FR4). More information can be obtained by clicking the respective info button (indicated by a question mark). Model parameters are grouped in terms of the tissue components to which they relate. Visual highlighting in the 3D alveolus model emphasizes these connections (FR5). For instance, all tissue components except the capillary are grayed out when the cursor is over the window for model parameters relating to the blood. To examine the process in the 3D model in more detail, it can be moved, rotated or zoomed. Detailed quantitative information can be obtained by hovering over a graph with the mouse. The instance menu allows direct comparison of different parameter settings by running several simulation instances simultaneously (FR6) (Figure 4, bottom). Characteristic coloring and custom naming facilitate distinguishing between different simulation instances. A selected instance can be copied, deleted or reset to its initial parameter values. Parameter presets for healthy and common pathogenic conditions are provided (FR6) (Table 2). Finally, the user interface contains control elements to monitor or reset simulation time and to toggle between pressure units and visual highlighting modes.

**Table 2:** Parameter value shifts in presets representing pathogenic conditions. For every condition, pathophysiological issues or symptoms are represented by increased (↑) or decreased (↓) values of the respective model parameters.

| Pathogenic Condition | Pathophysiology / Symptom | Parameter Value Shift |
| --- | --- | --- |
| Pneumonia | Fever<br>Tissue damage<br>Accumulation of fluids and dead cells | Temperature ↑<br>Surface area ↓<br>Barrier thickness ↑ |
| ARDS (acute respiratory distress syndrome) | Collapse (alveolar aelectasis)<br>Fever | Surface area ↓↓<br>Temperature ↑ |
| COPD (chronic obstructive pulmonary disease) | Impaired exhalation<br>Impaired exhalation<br>Tissue damage | Alveolar pCO<sub>2</sub> ↑ and blood pCO<sub>2</sub> ↑<br>Alveolar pO<sub>2</sub> ↓<br>Surface area ↓ |
| Pulmonary fibrosis | Thickened and scarred connective tissue<br>Impaired inhalation | Barrier thickness ↑<br>Alveolar pCO<sub>2</sub> ↓ |
| Pulmonary embolism | shunt<br>shunt | Blood volume ↓↓<br>Blood flow velocity ↓↓ |

### 3.4 Discussion on visualization and Interactivity

*Alvin* intends to increase understanding of the complex relationships of gas exchange by highlighting connections and allowing comparison of multiple simulations. Previous interactive systems for gas exchange also hold their great strength in this point. (Winkler et al., 1995) have modeled the lung as a complex of abstract gas exchange units (compartments) that can be simulated under individual conditions. (Kapitan, 2008) have created a model of gas exchange that is based on the alveolar gas equation (Sharma et al., 2020) and takes the ratio of ventilation to perfusion into account. Both systems enable simulation of inhomogeneous distribution of ventilation and perfusion. This provides valuable insights into higher-level relationships. In both systems, individual gas exchange units and the whole complex are visualized by means of abstract schematic representations. What happens in detail and how it looks like remains unanswered. *Alvin* fills this gap. The site of gas exchange is no longer abstract—a 3D model illustrates an alveolus in realistic proportions. It conveys the structure of important components (capillary net, tissue barrier). The connection between structure and function is interactively explored in the simulation. Blood flow and tissue thickness in the 3D model adapt to the parameter settings and directly affect the simulation process. What further sets *Alvin* apart from the two systems mentioned above is the possibility of running multiple simulation instances simultaneously. This allows different conditions to be compared directly instead of being modeled and explored one after the other.

The combination of providing parameter value presets as well as allowing parameter configurations by the user enables a presentation of the model that expands existing best-practice (Mogilner et al., 2011). *Alvin* includes a multitude of visualization elements and interaction possibilities. They aim at an intuitive usage of the application and understanding of the gas exchange simulation. It should be assessed whether the use of *Alvin* is actually perceived as intuitive. For this purpose, in the context of a use case study (described in Section 4.2), we had a group of users fill out a standardized questionnaire to measure intuitive usability.

## 4 Applying *Alvin*: Use Case Studies

We provide two concrete examples for the application of *Alvin*. First, we demonstrate how the interactive simulation can be used to interpret data from the literature. Second, we report on *Alvin*’s integration into a university level virtual class.

### 4.1 *Alvin* in Research: Interpreting Data and Testing Predictions

To present a possible use case of *Alvin* for research, we employ the application to check the plausibility of pulmonary diffusion capacity measurements. The pulmonary diffusion capacity 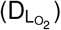 describes the lungs’ capacity to transport oxygen from the air to the blood. It is defined as the oxygen consumption 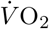 in L/min (oxygen uptake over time) divided by the mean oxygen pressure gradient between alveolar air and capillary blood ΔpO_2_ (Lindstedt, 1984).

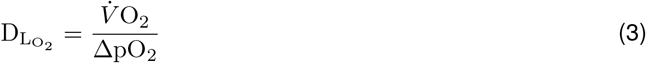

Physiological estimates of 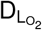 are usually derived from measurements of diffusion capacity for carbon monoxide 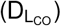 (Forster, 1964; Crapo and Crapo, 1983). Normal values of 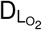 at rest are around 30 mL/(mmHg · min) (Hsia et al., 2016). Determination of 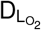 based on morphometric data has resulted in a value of 158 mL/(mmHg · min) (Weibel, 2009) and thereby exceeds physiological approximations considerably. There are several reasons for this discrepancy (Hsia et al., 2016). One of them is that for the morphological estimation, a complete perfusion of the capillaries is assumed and the entire alveolar surface is included in the calculations (Weibel, 1970). Under normal conditions, only about 50 % of capillary segments in the alveolar wall are perfused by erythrocytes and thus contribute to gas exchange (Okada et al., 1992) (Figure 5A). Increasing blood pressure (e.g. due to increased cardiac output) leads to recruitment of further capillary segments. In the perfusion fixed, air-filled lung, only about 60 % to 70 % of the alveolar surface area is exposed to air (Bachofen et al., 1987; Gil et al., 1979). In addition, lung volume changes during respiration depending on the transpulmonary pressure. It has been proposed that alveolar recruitment may be responsible for these volume changes, i.e., opening and closing of alveoli (Carney et al., 1999). However, *in situ* studies rather suggest an increase in alveolar size (D’Angelo, 1972). In terms of the model parameters in *Alvin*, both hypotheses manifest themselves in changes in the alveolar surface area available for gas exchange. A surface area of 207,000 µm^2^, measured in inflation-fixed lung tissue (Stone et al., 1992), describes a maximum surface exposure of 100 %. The default surface area setting in *Alvin* is 121,000 µm^2^ and thus corresponds to an exposure of 58 %. This value was taken from a study in which the tissue was perfusion fixed (Mercer et al., 1994). Capillary recruitment in *Alvin* is reflected in capillary blood volume, for which the default value 404,000 µm^3^ represents 50 % recruitment. By mimicking the ratios of capillary recruitment and alveolar surface area in *Alvin*, one can directly trace the effect on 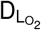. 100 % alveolar surface exposure and 100 % capillary recruitment in *Alvin* yield a 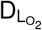 of 200 mL/(mmHg · min). 58 % alveolar surface exposure and 50 % capillary recruitment result in a 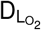 of 62 mL/(mmHg · min).

**Figure 5:**
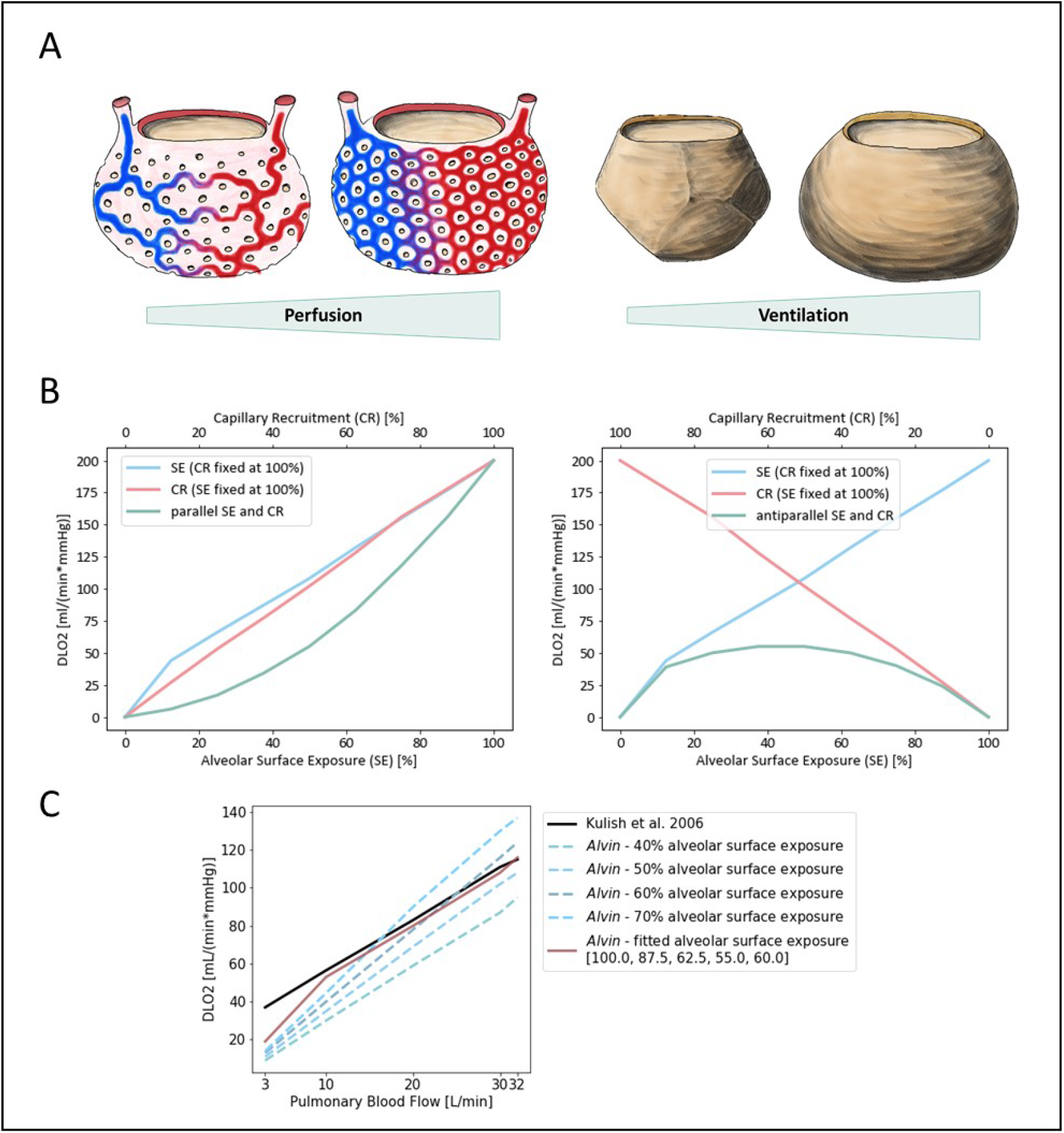
Diffusion capacity of the lung for oxygen 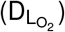 strongly depends on perfusion and ventilation. **(A)** Illustration of capillary recruitment (left) and alveolar expansion (right). **(B)** Diffusion capacity of the lung for oxygen 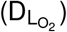 depending on capillary recruitment and alveolar expansion for a parallel (left) and antiparallel combination (right). Alveolar expansion and the ensuing surface exposure are simulated in *Alvin* by increasing alveolar surface area from 0 (0 %) to 207000 µm^2^ (100 %) in steps of 12.5 %. Capillary recruitment is represented by capillary blood volume increase from 0 (0 %) to 808000 µm^3^ (100 %) in steps of 12.5 % in *Alvin*. **(C)** Comparison to published 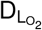 estimates (Kulish, 2006) (black). Pulmonary blood flow was interpreted as blood volume in *Alvin*, assuming a flow velocity of 1.5 mm/s and morphological features (mean capillary length of 500 µm (Weibel et al., 1993) and maximum volume of alveolar capillary bed 808000 µm^3^ (Ochs et al., 2004; Gehr et al., 1978)). Alveolar surface exposure was fixed at constant values (blue dashed lines) and adjusted with increasing pulmonary blood flow (red line).

Alveolar surface area and capillary recruitment impact 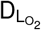 estimates almost linearly (Figure 5B). Additionally, it is interesting to observe their synergistic effect, as ventilation and perfusion are regulated to match (reviewed in (Wagner, 1981; Petersson and Glenny, 2014)). Parallel increase of both alveolar surface exposure and capillary recruitment lead to a non-linear increase in 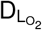, slowly at first and then more rapidly. Consistently, anti-parallel combination of these factors yields generally low 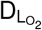 estimates, with a peak at 50 % each. Quantification of this relationship in *Alvin* can be used to interpret other data from the literature. For instance, 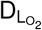 has been estimated from measurements of 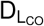 and pulmonary blood flow (Kulish, 2006). To recreate these estimates, pulmonary blood flow, expressed in volume per unit time, was interpreted as alveolar blood volume in *Alvin*. Assuming a constant blood flow velocity of 1.5 mm/s, the alveolar blood volume was obtained from the mean capillary length of 500 µm (Weibel et al., 1993) and the maximum volume of alveolar capillary bed 808,000 µm^3^ (Ochs et al., 2004; Gehr et al., 1978). Under these conditions, 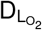 was determined in *Alvin* with varying alveolar surface area settings (Figure 5C). The resulting D_L_O2 graphs all differed in slope from the published data (Kulish, 2006). Thus, Kulish’s predictions did not appear to have been based on constant alveolar surface exposure. By adjusting alveolar surface area values (100, 87.5, 62.5, 55.0 and 60 % surface exposure) along with increasing blood flow (3, 10, 20, 30 and 32 L/min), the results could finally be reconstructed. This fitting was not successful at very low blood flow values.

This is only one example of how to employ *Alvin* to investigate correlations in a broader sense or to reproduce data from the literature to gain further insight. Further questions could address the kinetics of gas exchange. One possibility would be to investigate the threshold conditions under which the blood is still sufficiently oxygenated within the transit time.

### 4.2 *Alvin* in Higher Education: Physiology Lab Course

For the application in teaching, the benefits of an interactive simulation have been perceived and exploited early on (Dewhurst et al., 1988; Davis and Mark, 1990) and are still being pursued today (Jacob et al., 2012; Tworek et al., 2013). Therefore, we integrated *Alvin* into a university level class on human biology, specifically an online practical session on blood and respiration. *Alvin* was used to support the online session by providing an interactive model of the cooperation of the bloodstream and the respiratory system. The suitability of *Alvin* for this course was measured with an online questionnaire.

The course was scheduled for 2 hours and 45 minutes. The participants consisted of students of teaching Biology, specifically of the German levels of *Grundschule* (elementary school), *Mittelschule* (secondary school) and *Gymnasium* (grammar school). After an introduction into the topic “Blood and Respiration” in the form of a 45 minute lecture, *Alvin* was presented briefly, explaining how to use the application and interpret the 3D model and graphs. Participants were given a few minutes to familiarize themselves with *Alvin*. They were then asked for feedback as they worked with the application. An online questionnaire was provided to collect responses. Participation was voluntary and could be withdrawn throughout the event. Submitting the questionnaire as a whole, or answering individual questions, was not mandatory. The questionnaire was split in four parts. The entire questionnaire, translated from German, can be found in the supplementary material (Section S.2).

The first part consisted of a generic demographic questionnaire, extended by specific questions to assess the formal background of the students and their experience with the subject. We received N = 101 submission which were at least partially answered. Of the N = 101 surveys received, 11 self-identified as male, 81 as female. The second part contained 13 different exercises addressing respiratory processes in the alveolus. These exercises provided instructions on how to integrate *Alvin* into solution approaches. Among other things, these exercises highlighted well-known relationships and phenomena such as the Bohr effect (Riggs, 1988). The mean overall score for all participants was 2.07 on a scale of (1-4 with 1 indicating perfect answers). Missing answers were graded with the score 4. Most of the exercises were solved correctly. Few answers were incorrect, most score 4 ratings were due to missing answers.

The third part consisted of two standardized questionnaires to assess the visual aesthetics and the usability of the application: Visawi-s (Visual Aesthetics of Websites Inventory-short version) (Moshagen and Thielsch, 2021) and QUESI (Questionnaire for Measuring the Subjective Consequences of Intuitive Use) (Hurtienne and Naumann, 2010). Visawi-s (Moshagen and Thielsch, 2021) captures four central aspects of aesthetics from the user’s perspective: simplicity, diversity, colorfulness and craftsmanship. Participants were presented with statements targeting these four aspects. They rated them on a scale from 1 (strongly disagree) to 7 (strongly agree). The mean overall (N = 72) Visawi-s score was 5.8 (see Figure 6A). The standardized QUESI provided a measure of usability (Hurtienne and Naumann, 2010). It is based on the assumption that intuitive use is the unconscious application of prior knowledge leading to effective interaction. It can be divided into the following subscales: Subjective mental workload, perceived achievement of goals, perceived effort of learning, familiarity, and perceived error rate. The total score of the questionnaire is equal to the mean across all five subscales. Generally, higher scores represent a higher probability of intuitive use. Participants’ (N = 69) assessments of the use of *Alvin* resulted in a QUESI score of 2.98 (Figure 6A). Published benchmark values for mobile devices and applications (Naumann and Hurtienne, 2010) range from 2.39 (Alcatel One Touch 311) to 4.23 (Nintendo Wii). Familiar products generally perform better in the QUESI (Naumann and Hurtienne, 2010). Hence, participants’ prior experience with similar systems in a broader sense, for example, with computer games in general, is important. The majority of our participants (N = 59) reported rarely (yearly to never) playing computer games. The minority (N = 29) reported using computer games frequently (monthly to daily).

**Figure 6:**
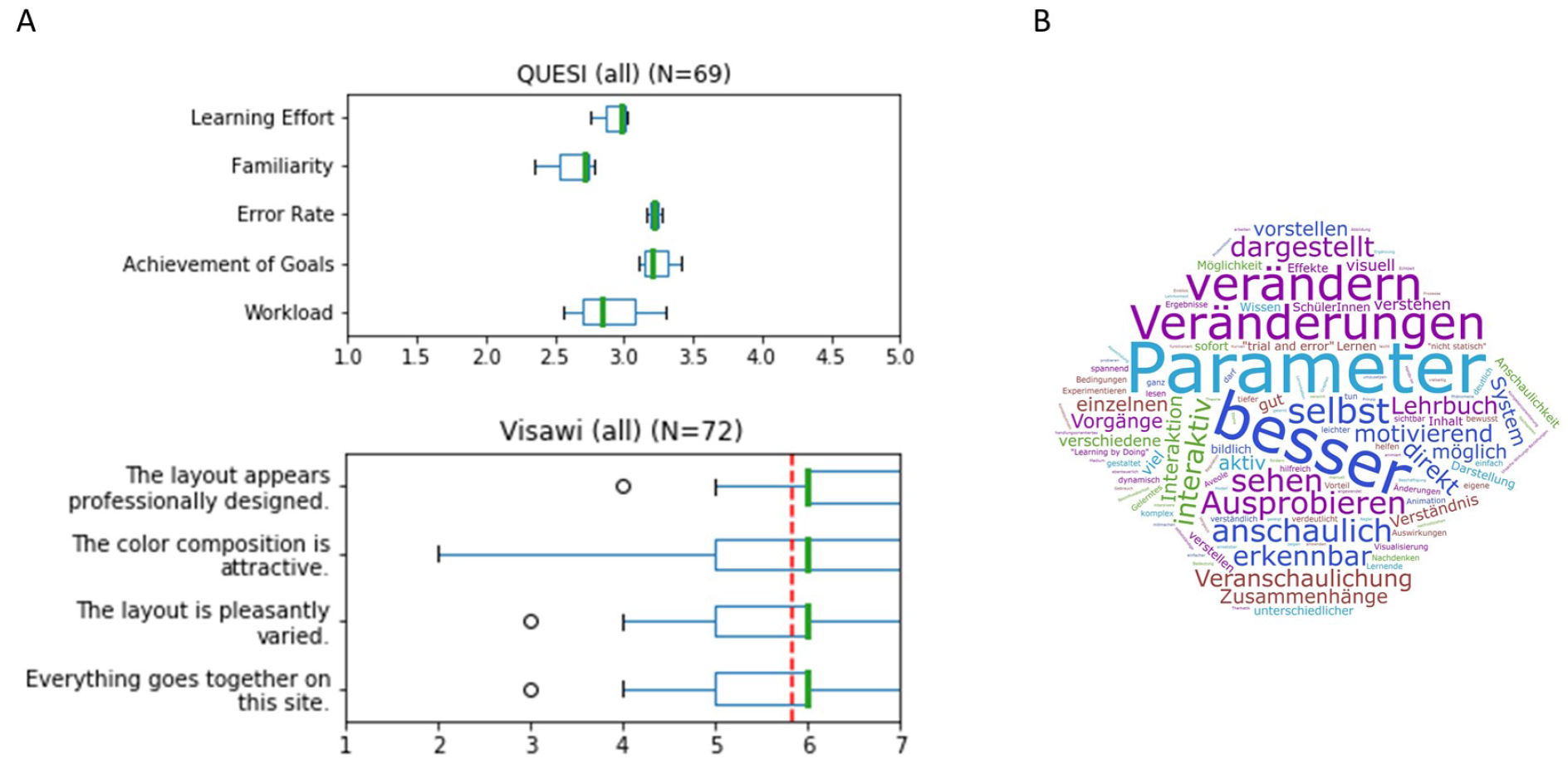
Results of a survey for undergraduate students that worked with *Alvin* in a physiology lab course. **(A)** Results on usability from the standardized survey QUESI (Hurtienne and Naumann, 2010) (top). Five subscales are assessed, with higher scores obtained the more intuitive the use of the system was perceived to be. The mean overall QUESI score from 69 forms was 2.98. The standardized Visawi-s survey (Moshagen and Thielsch, 2021) addresses design features. The 72 participants rated from 1 (strongly disagree) to 7 (strongly agree). The mean score over all four categories was 5.8 (red, dashed line). **(B)** Participants were asked “which benefits do you see in this system compared to a traditional text book?”. A frequency analysis on the answers was performed. The most recurrent terms were (translated from German): “parameter”, “better”, “modifiy”, “changes”, “by oneself”, “illustrative”, “testing”, “see”, “illustrated”, “apparent”, “interactive” and “immediate”.

Finally, the questionnaire included free-form questions aimed at the acceptance of the software in the educational context. One of them was “Which benefits do you see in this system compared to a traditional text book?”. A frequency analysis on answers revealed the highest recurrence for the terms “parameter”, “better”, “modifiy”, “changes”, “by oneself”, “illustrative”, “testing”, “see”, “illustrated”, “apparent”, “interactive” and “immediate” (Figure 6B). A question asking for general feedback was responded to in part with constructive criticism. In particular, it was noted that the content of *Alvin* and the subject-specific tasks were too complex for this introductory event. Or that more time would have been necessary to familiarize oneself with the application. In addition, some reported problems switching between the German lecture content and the English-language application. The participants solved the subject-specific exercises for the most part correctly. It can thus be concluded that *Alvin* is suitable to assist in solving such tasks. Responses to free-text questions suggest which aspects of working with *Alvin* stood out as particularly positive. These include the possibility to interact with the simulation by configuring model parameters and the freedom to independently test different conditions. It was also perceived positively that the simulated processes are presented very illustratively in *Alvin*.

### 4.3 Discussion of Use Cases

Our exemplary use cases show the applicability of *Alvin* in research and in education. We showed an investigation of the dependencies of 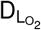 on surface area and blood flow in *Alvin*. Physiological estimates often only consider information about blood flow (Kulish, 2006). By reproducing these estimates in *Alvin*, one can draw conclusions about the alveolar surface. At particularly low blood flow values, it is not possible to reproduce the physiological estimates for 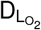 in *Alvin*. This could have different causes. In the logic of the model and the definition of 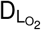, it is ensured that 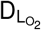 is zero when the blood volume is zero. The physiological estimates in (Kulish, 2006) do not seem to meet this criterion. (note: One cannot be certain, however, because in Kulish et al. (Kulish, 2006) the lowest reported value for blood flow is 3 L/min). It is possible that our model does not produce reliable results in the range of low blood volume values. Another possibility is that the derivation of 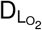 from 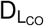 is not reliable in low ranges. This plausibility check shows how *Alvin* can be used to support or challenge published data. Drawing on known relationships, additional information can be obtained from previous results.

We also showed that *Alvin* is helpful for communicating respiratory processes in the training of undergraduate students. Well-known processes or phenomena like the Bohr-Effect (Riggs, 1988) can be recreated in *Alvin* and compared with results reported in the literature. Interactivity of the simulation enables experimentation with the model and exploration of its limitations. This aspect was also positively highlighted by participants of the physiology lab course in free-form answers of our questionnaire. The results of the QUESI and VISAWI questionnaires on their own do not allow for quantitative conclusions on usability or aesthetics of the application. This would require comparing them to corresponding results from comparable test situations (for example, about similar systems). At this point, one can only state that the replies did not hint at unknown issues. Instead, they were aligned with our expectations that participants should be able to operate the system autonomously and find its use appealing and relatively intuitive.

In summary, the integration of *Alvin* into physiology classes at the university level was successful. Beyond that, issues were pointed out where the implementation could be optimized in the future. Prominent and consistent were requests for more time to engage with *Alvin*. We deliberately refrained from providing the application to the participants in advance of this course to avoid a mutual influence of the participants regarding their experience with *Alvin*. This was important for the evaluation with the standardized questionnaires. For general use in teaching, however, this does not have to be taken into account. On the contrary, an exchange between students about the system could increase its learning value. We conclude that *Alvin* is less suitable to be included in a single physiology lesson. Instead, we recommend that students be made aware of the app ahead of time or to invest several course sessions.

## 5 Conclusion and Outlook

Interactive, visual simulations allow communicating modeling results and thereby help to further our understanding of the process under study. We presented *Alvin*, an application for simulating gas exchange in a single alveolus. The simulation is based on a mathematical model for the entire transport process of oxygen from the air to hemoglobin of the blood. We claim that having the goal of an interactive, visual simulation in mind when developing a mathematical model is beneficial for the modeling process. It resulted in a specific requirement for the model: In order to be able to map the course of the simulation on a three-dimensional tissue model, it had to be temporally and spatially resolved. Models evolve by being revised and improved over and over again (Drubin and Oster, 2010). If one assumes that a model can be better developed the more experts review it, then it is advantageous to make the model freely and intuitively accessible.

We argue that interactive visualization offers an engaging way to communicate theoretical models to other scientists and students. When cooperating with experimenters, it is important for theorists to present their models in the most accessible way possible. This creates as large a basis for discussion as possible in order to jointly plan further experiments or model refinements. By making model parameters intuitively configurable, any experimenter can compare his or her own measurements with the modeling results. By including undergraduate students in the target group for *Alvin*, we ensured that only a minimum of prior knowledge is required for its usage.

In the future, we plan to extend our model to encompass a system of multiple alveoli and their associated vessels. This will allow us to address further questions and complex relationships regarding gas exchange in lung tissue. It is known that the ventilation-perfusion relationship, and therefore the diffusion-perfusion relationship, has a strong influence on 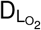 (Hyde et al., 1967; Hammond and Hempleman, 1987). An evolution of *Alvin* that includes an alveolar sac or a whole acinus with differently ventilated and perfused alveoli can provide valuable insights. This could also be used, for example, to further investigate the hypothesis of precapillary oxygen uptake (Tabuchi et al., 2013). It states that the oxygenation process already takes place in the precapillary arterioles before the blood reaches the alveolar capillary bed.

Rather than just presenting the data that results from a newly developed model, it is worthwhile to implement the model in a way that allows for interaction. Visualizing the simulation makes the engagement with the model more intuitive and accessible to a broader target group. Empiricists and theorists look at a system from different angles. Some work in a bottom-up fashion and take local samples and draw conclusions for the overall system. Others create abstract models for the overall system top-down and try to approach the truth by introducing more and more details. Only by working closely together can these two perspectives efficiently contribute to reliable results and become a “middle-out” approach (Noble, 2008). The communication of the achieved findings or predictions plays an important role here. We contend that interactive, visual simulations of theoretical models, as we have implemented with *Alvin* on respiratory processes in the alveolus, will make an important contribution to bridging the gap between empiricists and theorists.

## Author Contributions

*Alvin* concept and design: AK, AM, KS. Implementation of *Alvin*: AK, AM, KS. Model development and validation: KS. Planning and supervision of the use case in teaching: KP, AK, KS. Demonstration of possible application in research: KS. Supervision: SF, SvM, KP. Manuscript preparation: KS, SF, AK, SvM. All authors contributed to the article and approved the submitted version.

## Acknowledgments

We thank Andreas Hocke and Katja Hönzke for inspiring discussions and support of the project. We thank Wolfgang Kübler and Matthias Ochs for valuable feedback on *Alvin* and members of the CCTB for testing *Alvin*.

## Supplementary Material

### S.1 Methods

#### S.1.1 Implementation of *Alvin*

The application *Alvin* was built in Unity version 2020.1.16f1 (https://unity.com/). Unity is a platform for creating interactive real-time content. The mathematical model (see Section 2.1) provides the basis for the simulations. Parameter value changes in *Alvin* result in instantaneous updates of the visual output. All calculations are performed in the pressure unit mmHg. For the purpose of visualization, simulation time is slowed down by a factor of 40 compared to the gas exchange process *in vivo*.

#### S.1.2 Three-dimensional visual model of an alveolus

The three-dimensional, mesh-based geometric model of an alveolus was created in Blender^®^ version 2.82 (https://www.blender.org/). The model resembles a real alveolus not only in its general appearance, but also in proportions. The tissue barrier, consisting of alveolar lining fluid, epithelial cells and connective tissue fibers are represented by a single layer in this model. This layer forms a truncated sphere with diameter, volume and surface area comparable to the corresponding values of an alveolus reported in the literature (Table S.1). A network of hollow channels was modeled around this tissue layer, representing the capillary network enveloping the alveolus. The properties of the model network are chosen such that the relative magnitude of the volume and the surface area, as well as the relative radius and the length of the individual segments agree with the respective measured values from the literature (Table S.1). To facilitate visualisation of the blood flow, a capillary was cut open longitudinally, thereby exposing the inside of the channel. In addition, the inflow and outflow of blood is indicated with the help of additional capillaries that connect to the capillary net.

**Table S.1:** The visual three-dimensional model of an alveolus was created in Blender^®^. The size ratios were based on morphometric values from the literature.

| Parameter | Value from literature | Reference | Blender model |
| --- | --- | --- | --- |
| Alveolar diameter [$\mu\text{m}$] | 225 | (Mercer et al., 1994) | 225 |
| Alveolar volume [$\mu\text{m}^3$] | $4.2 \cdot 10^6$ | (Ochs et al., 2004) | $5.4 \cdot 10^6$ |
| Alveolar surface area [$\mu\text{m}^2$] | 121000 | (Mercer et al., 1994) | 150000 |
| Capillary volume [$\mu\text{m}^3$] | 808000 | abstracted from (Gehr et al., 1978; Ochs et al., 2004) | 787000 |
| Capillary surface area [$\mu\text{m}^2$] | 479000 | abstracted from (Gehr et al., 1978; Ochs et al., 2004) | 335000 |
| Capillary radius [$\mu\text{m}$] | 3.15 | (Mühlfeld et al., 2010) | 4.28 |
| Capillary segment length [$\mu\text{m}$] | 5.92 | (Mühlfeld et al., 2010) | 8.62 |

#### S.1.3 Simulation Output

In the graph panel, dynamic plots record the course of the simulation quantitatively. The plot “oxygen saturation along capillary” presents results obtained directly from the simulation calculations. The oxygen uptake, presented in another graph, is calculated assuming a standard amount of 270 · 10^6^ hemoglobin molecules per erythrocyte (Pierigè et al., 2008). Considering the parameter values from the configuration menu, but independent of the rest of the simulation, the “oxygen dissociation curve” is calculated for a range of partial pressure values of oxygen.

If several simulation instances are active at the same time, the respective results are displayed together in the graphs, while only information about the selected instance is considered in the 3D visualisation in the center.

Two different prototypes of *Alvin* were implemented for the different use cases described in this work. One prototype was adapted for educational use (Section 4.2) such that the application has two levels of different complexity. In the first level, only one simulation instance can run at a time. The second level with full complexity is unlocked by an access code. The other prototype features additional readouts. “Membrane” diffusing capacity for oxygen DMO_2_ and reaction half-time were required for model validation (Section 2.2). For the application example in research (Section 4.1), the output for diffusion capacity of the lung for oxygen was added. This second prototype is depicted in Figure 4 available for download at https://go.uniwue.de/alvin.

### S.2 Questionnaire for Evaluation of the integration of *Alvin* in a university level physiology lab course

The questionnaire was translated from the German original.

#### Demographics

In this section, we ask you to answer questions for general demographic information. These are relevant for a correct interpretation of your further answers.

1. Please indicate your age.
2. Please indicate your biological sex.
3. Do you have a visual impairment and will it be compensated for while using the system?
4. Are you affected by color vision deficiency or color blindness?
5. What handedness do you have?
6. On average, how often do you use the following media?
  - Internet
  - Computer (in general)
  - Computer games
  - Smartphone
  - Tablet
7. How would you rate your fluency in German? In this section, we ask you to answer questions about your prior knowledge in the subject area of the course and regarding your previous educational background.
8. In the context of which study program are you attending this event?
9. What semester are you in?
10. Did you attend the Human Biology lecture in the summer semester of 2020?
11. Have you studied the literature recommended in the above lecture on the subject of respiration?
  - N.A. Campbell and J.B. Reece. *Biology*. Always learning. Pearson Deutschland, 2015. ISBN: 9783868942590.
  - Robert F. Schmidt, Florian Lang, and Manfred Heckmann. *Physiologie des Menschen*. Springer-Lehrbuch. Springer-Verlag Berlin Heidelberg, 2011. ISBN: 978-3-642-01651-6.
12. Do you have other relevant prior knowledge from other sources?
  - School
  - Apprenticeship
  - Personal initiative

#### Subject-related exercises

In this group of questions, you will be given tasks that you can answer using the system. We ask you to discuss comments on the use of the app only in a joint round at the end of the event in order to minimize influencing the other participants. Now, familiarize yourself with the application. Look at how the graphs change in response to the controllers. Also observe how different disease patterns affect the values.

1. Which correlations between the course of the oxygen saturation curve (“Oxygen saturation along capillary”) and the visualized simulation can you identify? In this and the following blocks of questions, you will be given tasks to answer using the system. After each task (there are 3 tasks in total, each with subtasks), the answers will be discussed in plenary. We ask you to discuss comments on the use of the application only in a joint round at the end of the event in order to minimize influencing the other participants.
2. How does the oxygen dissociation curve change, when the body temperature rises to 40 °C (fever)?
3. How does this affect the ability of hemoglobin to bind oxygen in the lungs?
4. How does it affect the ability of hemoglobin to deliver oxygen to tissues? Fever is normally accompanied by an increase in respiratory rate. By increasing the respiratory rate, the increased CO_2_ produced by the increased metabolism during fever can be better exhaled. The partial pressure of CO_2_ in the blood affects the ability of hemoglobin to bind O_2_.
5. By how many mmHg must the venous CO_2_ partial pressure be lowered to achieve the same oxygen saturation at 40°C as at 37°? The cruising altitude of passenger aircrafts is around 10 to 13 km. At this altitude, the partial pressure of oxygen is only between 30 and 44 mmHg. Therefore, the air pressure in the cabins of passenger aircrafts is artificially increased, but only to a level corresponding to the air pressure at about 2000-2500 m above sea level. Thus, an oxygen partial pressure of approx. 60 mmHg is achieved in venous blood.
6. What oxygen saturation does this correspond to?
7. At what alveolar pO_2_ can a healthy person achieve this?
8. What oxygen saturation does the blood of a patient suffering from COPD reach at the same atmospheric pressure?
9. What happens to the oxygen saturation of a person suffering from COPD if he or she develops a fever during a flight? In this block of questions, you will be given tasks to answer using the system. Please configure the application using the activation code provided in the lecture. We ask that you do not discuss comments on the use of the application until a joint round at the end of the course to minimize influencing the other participants.
10. Sometimes a lung has to be surgically removed due to a disease. What effects does this have on the oxygen saturation of the blood? In tissues with very high metabolic rates, for example heavily used muscles, the CO_2_ concentration can increase.
11. How does this affect the oxygen dissociation curve?
12. How does it affect oxygen uptake in the lungs and oxygen delivery in the tissues? Start two simulation instances with the parameters for a healthy person. Increase the partial pressure of CO_2_ in the arterial blood of one instance to 75 mmHg. Using the oxygen dissociation curves, measure the absorbed oxygen in the lungs and the delivered oxygen in the tissues.
13. At which blood pCO_2_ is more oxygen available to the tissue? This phenomenon is called the Bohr effect. Athletes, especially high-altitude mountaineers, can adapt to conditions at high altitude by training for longer periods at low oxygen partial pressure. This increases the diphosphoglycerate (DPG) concentration in the erythrocytes. Now, we would like to understand why this is beneficial. Start a simulation instance with the parameters for a healthy subject. First, reconstruct the conditions that exist when climbing at high altitudes: Decrease atmospheric pressure until alveolar pO_2_ drops to a low value such as 40 mmHg. Arterial pO_2_ is also reduced in these conditions. Set this to 30 mmHg.
14. What is the oxygen saturation of the blood? Duplicate the instance. Now, set the DPG concentration in one of the two instances to the maximum value (adjustment to high altitude).
15. What happens to the oxygen dissociation curve?
16. What happens to the oxygen saturation of the blood?
17. The effect you observed initially appears to be rather disadvantageous. Now, measure the oxygen saturation in the lungs and in the tissue in the respective graphs and determine the difference between these values.

The questions will now be discussed in the plenum of the event. We ask you to discuss comments on the use of the application only in a joint round at the end of the event to minimize influencing the other participants.

The phase of active use of the system is now complete. In the following, we ask you to answer questions about your user experience. This is for systematic evaluation of the system. Please note that for the first two questions a “soft” inquiry will appear if you do not answer them or answer them only partially. The corresponding groups of questions are standardized questionnaires, where a complete answer has a lot of value. Of course, you can still skip them unanswered if you wish.

#### QUESI – Questionnaire for Measuring the Subjective Consequences of Intuitive Use

This is a standardized questionnare (Hurtienne and Naumann, 2010). Try to base your assessment of the system solely on the use of the system (and not, for example, on the difficulty of the task itself). There are no right or wrong answers. Please answer spontaneously and do not omit any questions.

Answer scale with equidistant levels: 1 = “Fully disagree”, 2 = “Mainly disagree”, 3 = “Neutral”, 4 = “Mainly agree”, 5 = “Fully agree”.

1. I could use the system without thinking about it.
2. I achieved what I wanted to achieve with the system.
3. The way the system worked was immediately clear to me.
4. I could interact with the system in a way that seemed familiar to me.
5. No problems occurred when I used the system.
6. The system was not complicated to use.
7. I was able to achieve my goals in the way I had imagined to.
8. The system was easy to use from the start.
9. It was always clear to me what I had to do to use the system.
10. The process of using the system went smoothly.
11. I barely had to concentrate on using the system.
12. The system helped me to completely achieve my goals.
13. How the system is used was clear to me straight away.
14. I automatically did the right thing to achieve my goals.

#### Visawi-s - Visual Aesthetics of Websites Inventory-short version

This is a standardized questionnare (Moshagen and Thielsch, 2021). On a scale of 1 (strongly disagree) to 7 (strongly agree), please rate the extent to which you agree with the following statements regarding the system.

1. The layout appears too dense. (r)
2. The layout is pleasantly varied.
3. The color composition is attractive.
4. The layout appears professionally designed.
5. The layout is easy to grasp.
6. The layout is inventive.
7. The colors do not match. (r)
8. The layout is not up-to-date. (r)
9. Everything goes together on this site.
10. The design appears uninspired. (r)
11. The choice of colors is botched. (r)
12. The site is designed with care.
13. The site appears patchy. (r)
14. The layout appears dynamic.
15. The colors are appealing.
16. The design of the site lacks a concept. (r)
17. The layout appears well structured.
18. The design is uninteresting. (r)

Negatively-keyed items are indicated by (r) and are reverse-scored.

#### Customized questions on the use of *Alvin*

1. On a scale of 1 (strongly disagree) to 7 (strongly agree), please rate the extent to which you agree with the following statements regarding the system.
  - I frequently changed the “Incoming Deoxygenated Blood” parameter values for completing the tasks.
  - I frequently changed the “Alveolar Space” parameter values for completing the tasks.
  - I frequently changed the “Tissue Structure” parameter values for completing the tasks.
  - The system supported me in the configuration and interpretation of the parameters.
  - I was confused by the information provided by the system.
  - Assessment of the parameter values was useful for my understanding of the processes.
  - I found the ability to create, configure, and compare multiple instances useful.
  - I found the ability to copy instances useful.
  - I found the ability to reset instances to initial configuration useful.
  - I have used the output graphs frequently during my use of the system.
  - I could easily extract the information relevant to me from the graphs.
  - I regularly read the exact numerical values of a plot using the mouse-over function.
  - I found the visual highlighting of the simulated components of the alveolus when the cursor was over a parameter group helpful.
  - I found the visual highlighting distracting.
  - I have disabled visual highlighting for most of the time I used it.
  - I found the ability to reset the simulation time helpful.
  - The mouse-over tooltips help assisted me in using the system.
2. On which device or which version(s) of the system did you use? You can use detailed information and multiple selections if, for example, you used multiple usage paths. Detailed information about the operating system (for example, “Windows 10 version 1903”, “macOS 10.13”), as well as the device (for example, processor (i5-5700) or graphics card (GeForce 2 MX), or computer model (MacBookPro Late 2015)) or browser (for example, Firefox 83.0, Safari 12) is helpful, especially if problems occurred.
  - Windows Desktop (.exe)
  - macOS Desktop (.app)
  - Linux Desktop
  - Browser (WebGL)
  - iOS Tablet
  - Android Tablet
  - iOS Smartphone
  - Android Smartphone
3. On a scale of 1 (strongly disagree) to 7 (strongly agree), please rate the extent to which you agree with the following statements regarding the system.
  - The system responded to my input immediately.
  - Animations were smooth and without annoying leaps.
  - The performance of the system affected my desired use.
4. Which benefits do you see in this system compared to a traditional text book?
5. For which topics from your previous studies would you have appreciated a comparable application?
6. Assuming you’ll be teaching physiology - Could you imagine integrating this application into your own teaching?
7. Could you imagine using a similar system on an appropriate topic in your classes (or a similar event)?
8. Please share general comments, suggestions and feedback.

